# Effects of Fermented *Idesia Polycarpa* Residues supplementation on laying performance of Laying Quails

**DOI:** 10.1101/471201

**Authors:** Xinanbei Liu, Na Li, Yueyue Shu, Yiran Sun, Yu Li, Wang Hua, Yang Ye, Fang Chen, Lin Tang

## Abstract

Although *Idesia polycarpa* oil has been wildly explored as a raw material for biodiesel, the reports studying the by-product *Idesia polycarpa* fruit residues (IPR) are few. This study aimed to evaluate the effect of the *Idesia polycarpa* fruit residues fermented feed additive (IPFF) on the egg production of laying quails. The egg production and related performances include egg quality, yolk cholesterol, yolk fatty acid, quails’ jejunum morphology, and relative gene expression were determined in this study. Compared to the standard diet group, birds fed the 1% IPFF showed a higher egg production (87.7% on average, 11.5% above the control; P<0.01). The yolk fatty acid composition and n6/n3 ratio were affected by IPFF or IPR. Compared to the standard diet group, the egg cholesterol content was lower in both IPFF and IPR groups, and the yolk n6/n3 ratio in the 5% IPFF group (10.3; P<0.01) was more reasonable. Meanwhile, birds under IPFF dietary supplement showed a thicker jejunum wall, higher villus, and deeper crypt than the standard diet group. In addition, the altered mRNA expression of four genes involved in cholesterol and fatty acids metabolism (SREBP-1, SREBP-2, ADGL, APOVLDL-II) in the 1% IPFF group and 5% IPR group indicated that the lipids metabolism and transportation were enhanced in the interclavicular fat pad and liver, relative to the standard diet group.

**HIGHLIGHTS:** Egg production was higher in IPFF groups

Egg cholesterol was lower in IPFF groups

Lipid metabolism and transportation was enhanced in IPFF groups

Intestine wall was thicker in IPFF groups

## 1. Introduction

*Idesia polycarpa* (*Idesia polycarpa* Maxim. var. *vestita Diels*) is wildly distributed in the south of China. *Idesia polycarpa* oil is edible and also explored as a raw material for biodiesel. (Yang F-X et al. 2009; Wang QY et al. 2013). IPR is the byproduct of the oil squeeze procedure which contains many bioactive substances. Extractions of IPR showed various bioactivities include remarkable antioxidant ability, anti-skin-aging ability, and anti-adipogenic ability in previous studies. Indicated that IPR has the potential for natural product extraction or animal feed exploitation.

From a historical perspective, the phytogenic fermented feed was recognized to benefit the farm animals and described as far back as the beginning of the last century (Curtis 1909; Thom and Church 1921). Fermentation could effectively enhance the nutrition of residues with the decrease of anti-nutritive factors and allergenic proteins (Emenalom et al. 2012; Li C-Y et al. 2014; Dwivedi et al. 2015). Besides that, fermented feed is commonly regarded as a plausible alternative to traditional antibiotics in the cultivation industry (Hong et al. 2004; Karásková et al. 2016). Hence, fermented feed is still gaining interest as it plays an essential role in the quality of feed production and the utilization of unexploited materials as one of the most common materials for fermentation in feed researches. (Zhang et al. 2020).

Quail egg is known for its good taste, health benefits, and Nutrient-rich. (Aba et al. 2016; Ibukun and Oladipo 2016). Besides that, quail eggs of *Coturnix coturnix japonica* have often been used in animal research (Chah et al. 1976) as a sensitive experimental animal (Cardoso et al. 2011; Laurence et al. 2014). It also was a appropriate choice for testify the effect of fermented *Idesia polycarpa* products on the quail’s egg production. As we know, fatty acid and cholesterol metabolism are involved in the regulation of egg formation (Speake et al. 1998; Wu et al. 2013; Chen et al. 2016). There was evidence that extractions in IPR evolved in the lipids metabolism(Lee et al. 2013). That means IPR or IPFF have the potential to enhance the egg production of laying quails by altering the lipid metabolism. Therefore, the present study aims to testify the performance of IPFF on the production of laying quails as a feeding stuff additive.

## 2. Materials and methods

### 2.1. *Idesia polycarpa* fermented feed

Defatted oil residues of *Idesia polycarpa* obtained by the physical cold pressing process were kindly supplied by the Agriculture Bureau of Jianyang, China. IPFF was produced in our laboratory using the solid fermentation method by *Candida utilis* (purchased from China Center of Industrial Culture Collection, CICC), and the basal feed purchased in the local market contains all nutrients necessary for animal feeding which was also supplied for the local layer quail farms. Nutrients of IPFF, IPR, and basal diet (BD) are shown in **Table 1**.

**Table 1.**
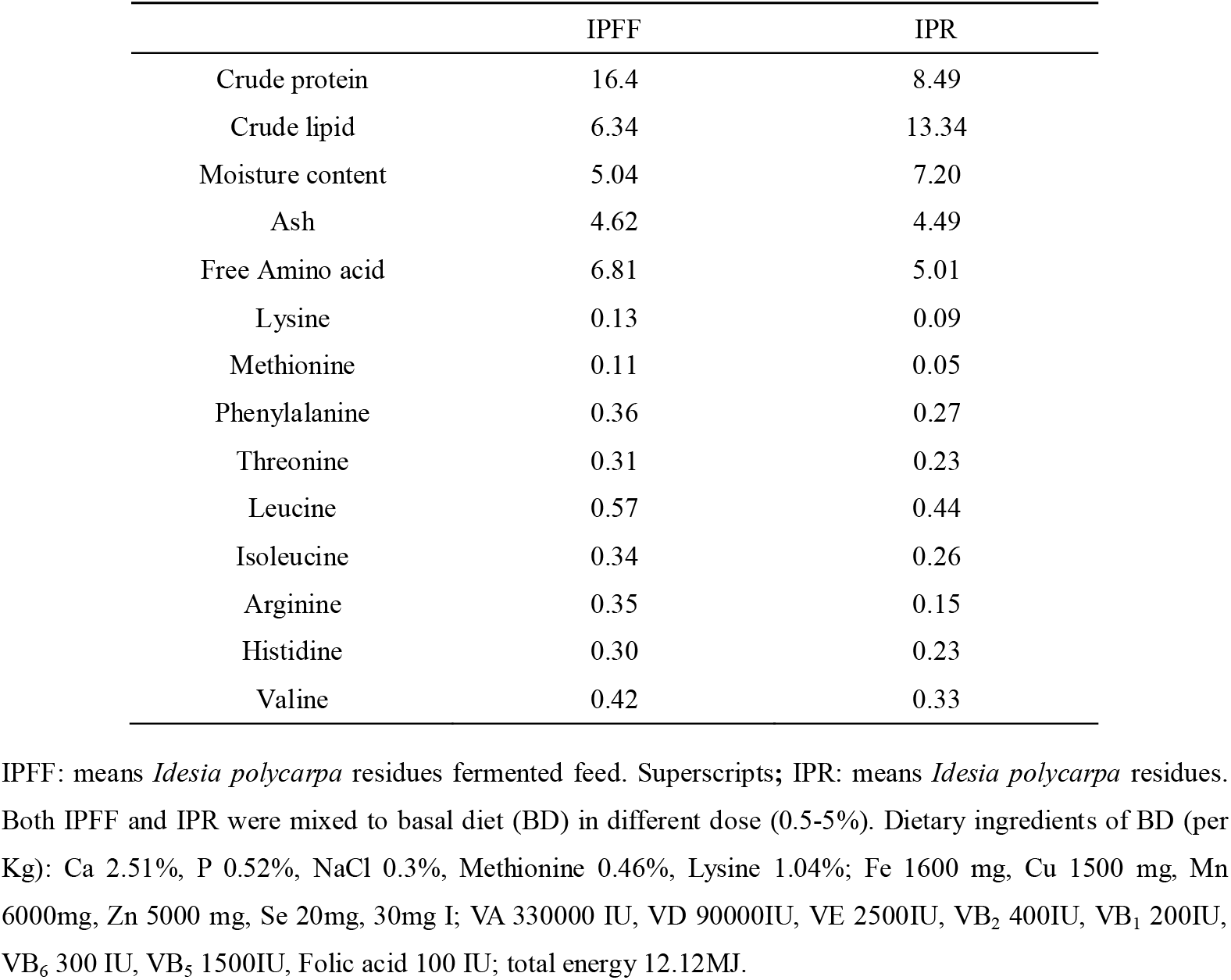
The Main Nutrients and Amino acids of IPFF and IPR (%)

### 2.2. Experimental animals and feeding management

All animal protocols were approved by the Sichuan University Animal Care and Use Committee. A total of 300 laying quails (*Coturnix coturnix japonica*, 90 days old) from a local breeding farm (Chengdu, China) were used. All birds were kept in a room with a relatively stable temperature (24-30 °C) and humidity (65%), and the fed followed the standard procedure (All the feeding conditions were as same as the commercial birds which were feeding in the farm). Those birds were randomly allocated into 6 groups (50 quails in each group; Control, base diet only; A, a basal diet supplemented with 5% IPFF; B, a basal diet supplemented with 1% IPFF; C, a basal diet supplemented with 0.5% IPFF; D, a basal diet supplemented with 5% IPR; E, a basal diet supplemented with 1% IPR) according to the IPR or IPFF additive amount. During the 10 weeks experimental period, birds were fed with a fixed amount of diet (1000 g/d, which contained adequate nutrition such as protein, based on the farm feeding management for this season, equivalent to a dose of 20 g/d/bird), but they had access to water *ad libitum*. The quail’s differentiated diet was supplied during the 6 weeks feeding test period from week 3 to week 8 (Phase 2), and all groups were fed with basal diet only in the final three weeks (week 9-11; Phase 3). The feeding experimental design is shown in **Table 2**.

**Table 2.** Daily Diet Arrangement of IPFF, IPR, Control Groups in This Study (g)

|  | Week 0 to 2 |  |  | Week 3 to 8 |  |  | Week 9 to 11 |  |  |
| --- | --- | --- | --- | --- | --- | --- | --- | --- | --- |
|  | BD | IPFF | IPR | BD | IPFF | IPR | BD | IPFF | IPR |
| A | 1000 | - | - | 950 | 50 | - | 1000 | - | - |
| B | 1000 | - | - | 990 | 10 | - | 1000 | - | - |
| C | 1000 | - | - | 995 | 5 | - | 1000 | - | - |
| D | 1000 | - | - | 950 | - | 50 | 1000 | - | - |
| E | 1000 | - | - | 990 | - | 10 | 1000 | - | - |
| Control | 1000 | - | - | 1000 | - | - | 1000 | - | - |

BD: means basal diet; IPFF: means *Idesia polycarpa* residues fermented feed. Superscripts; IPR: means *Idesia polycarpa* residues. (20 g per bird)

### 2.3. Sample collection

The egg number was calculated every day, and the results were represented in the daily egg number as the mean value of each week. Eggs were randomly collected from each group (5 eggs from each group, 30 eggs in total) each week to measure the egg quality during the experimental period. Egg weights, density, axis ratio, egg yolk weight, and the ratio between egg weight and egg yolk weight were measured. The Haugh unit values were calculated using the Haugh unit formula, based on egg weight and albumen height, as determined by a height vernier caliper. Eggshell thickness and membrane were determined as the mean value of measurements taken at three locations on the egg (sharp end, blunt end, and middle section).

The egg yolk samples were separated from the rest of the egg for triglyceride and cholesterol determination at week 6 to 8. Yolk triglyceride and cholesterol were determined by ELISA (SMP500-15859-SBRE, Molecular Devices, America) using commercial kits, according to the manufacturer’s protocol (Jiancheng, Nanjing, China). At week 8, 3 additional eggs per group were collected for yolk lipid extraction.

At the end of Phase 2 (week 8), 3 quails of each group were randomly sacrificed by cervical dislocation for sample collection after being weighed. Tissues including intestine, interclavicular fat pad, and liver samples were immediately collected from all the 18 quails after their sacrifice. The same segment of the intestine was collected and stored in the prepared stationary phase (10% formalin) for histology analysis, fat pads were quickly weighted, both fat pads and liver samples were snap-frozen in liquid nitrogen and stored at −80 °C until the subsequent RNA isolation.

### 2.4. Yolk lipid extraction

Yolk samples were separated from boiled eggs. Yolk lipid was extracted (three yolks per group) by hexane extraction (AOAC 2003. 06). The extracted yolk lipids were stored at −20 °C until the subsequent fatty acids analysis.

### 2.5. Methylation and GC/MS

The methylation of yolk lipid was determined using the alkali catalysis method (Ulberth 1992). Peak area and fatty acid percentages were calculated using GC/MS-solution (Shimadzu, Japan), and fatty acid methyl esters were expressed as a percentage of total fatty acids.

### 2.6. Jejunum morphology

After being fixed in 10% formalin for 24 h, cross-sections of the jejunum tissues were selected for 5 μm thick paraffin section and stained with hematoxylin-eosin (HE). The analysis included the wall thickness, villus height, and crypt depth was performed by a microscope equipped with a camera and connected to a computer with the appropriate software (Olympus, BX53, software Cell B, Japan). Data were collected from at least five random visible regions under the microscope. The mean values (three birds per group) were calculated for subsequent statistical analysis.

### 2.7. RNA isolation

Total RNA was extracted from the fat pad and liver tissue using TRIZOL reagent (Takara, USA). The RNA quality and quantity were determined using nanodrop 2000 (Thermo Scientific, USA). The absorption (260/280nm) ratios of all extracted samples were between 1.8 and 2.0, and RNA integrity was further analyzed by gel electrophoresis. The first-strand cDNA was immediately synthesized using the PrimeScript RT reagent kit (Takara, USA). The obtained cDNA was stored at −20 °C until the subsequent real-time PCR analysis.

### 2.8. qRT-PCR analysis

Primers used for target genes amplification were either designed using primer 3.0 based on quail-related gene sequences or reported by specific reference (**Table 3**). *β*-actin was selected as a housekeeping gene to normalize target gene expression. All primers in this study were synthesized by TSINGKE biotech (Chengdu, China). The specificity of each of the designed primers was checked via gel electrophoresis analysis and melting curve analysis during quantitative real-time PCR. Relative quantification of all transcripts was performed by qRT-PCR using the Bio-Rad CFX96 real-time PCR system (Bio-Rad Laboratories, USA). Real-time quantitative PCR was performed with SsoFast EvaGreen Supermix (Bio-Rad Laboratories, USA). A total volume of 10 μl containing 5 μl of SsoFast EvaGreen Supermix, 3 μl of RNA free water, 0.5 μl of forward primer, 0.5 μl of reverse primer, 1 μl of template cDNA was used. PCR amplification was carried out as follows: denaturation at 95 °C for 30 s, followed by 40 cycles at 95 °C for 5 s, specific annealing temperature at 55 °C for 30 s. The 2^−ΔΔCT^ method was used to evaluate mRNA expression levels (Livak and Schmittgen 2001).

**Table 3.**
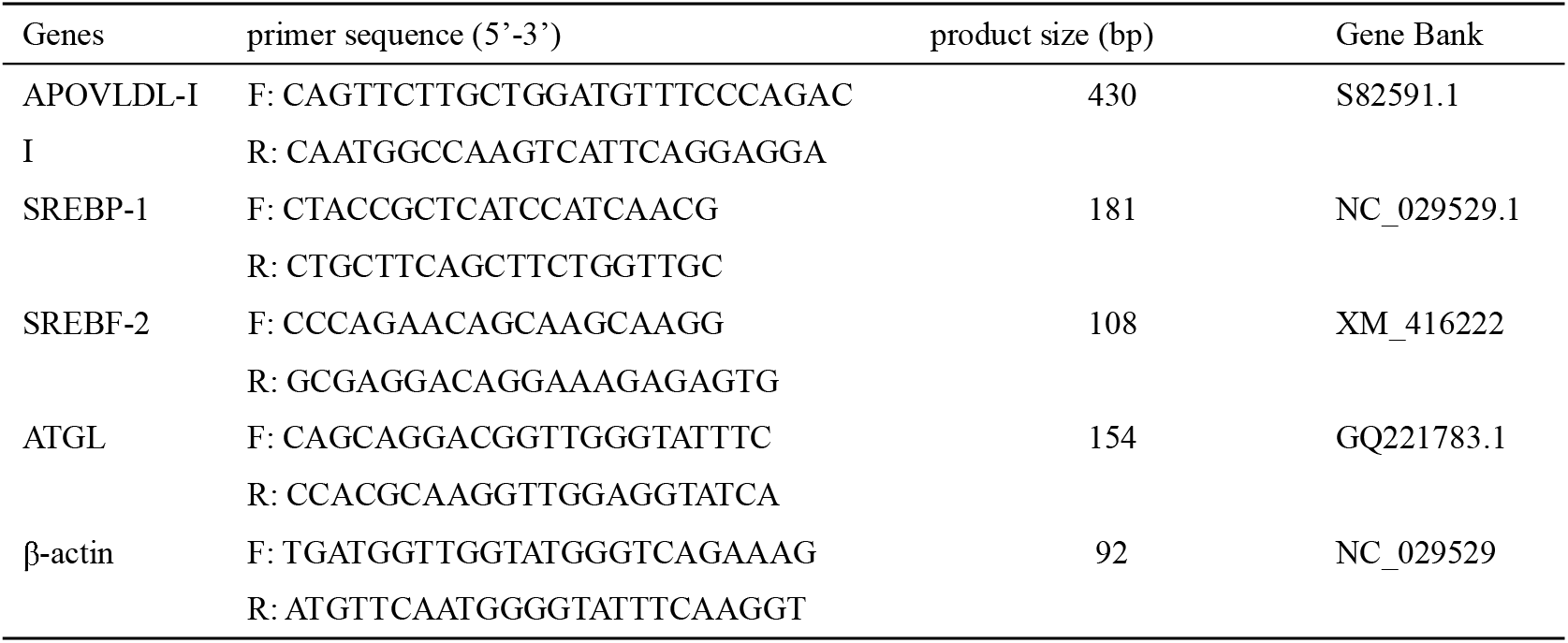
Primer Sequences for the mRNA Expression Analysis of Genes in This Study.

### 2.9. Statistical analysis

Statistical analysis was measured by using the SPSS software (SPSS, IBM, version 20). The standard error of means (SEM) obtained from the analysis of variance (ANOVA) that was generated. The significance of difference among means was determined by using one-way ANOVA and compared by Dunnett’s multiple range tests. The results were considered statistical significant at *P*< 0.05.

## 3. Results and discussion

### 3.1. Genes

Adipose triglyceride lipase (ATGL) is highly expressed in adipose tissue and catalyzes the initial step of triglyceride hydrolysis (Zimmermann and Zechner 2004).In the study of Chen et al, G0/G1 switch gene 2 (G0S2) overexpression carried out the inhibition of ATGL expression and resulted in a delayed laying onset and reduced egg production (Chen et al. 2016). Sterol regulatory element-binding protein-1 (SREBP-1) and SREBP-2 are SREBPs encoding genes that mainly regulate the homeostasis of lipid and cholesterol, respectively (Brown and Goldstein 1997; Eberlé et al. 2004). SREBP-1 up-regulation preferentially enhances fatty acid and triglyceride synthesis, while an increased SREBP-2 expression promotes cholesterol synthesis (Li X et al. 2014; Dong et al. 2015; Park et al. 2016). Apo Very Low Density Lipoprotein II (APOVLDL-II) is related to the secretion of VLDL, which plays a pivotal role in egg formation (Yang S et al. 2013). The described change of fat pad and the ATGL expression in the research of Chen et al (Chen et al. 2016) was also observed in our study (**Fig. 1**, **Fig. 2**). In group B, as compared to control, SREBP-1 was up-regulated in the fat pad (*P*<0.05) but was not significantly changed in the liver, while SREBP-2 was significantly down-regulated in the fat pad and up-regulated in the liver, indicated an enhanced lipids synthesis in the fat pad and enhanced cholesterol synthesis in the liver. Furthermore, the significant up-regulation of APOVLDL-II indicated that the secretion of VLDL was enhanced in both groups B and D (**Fig. 2**, P<0.01).

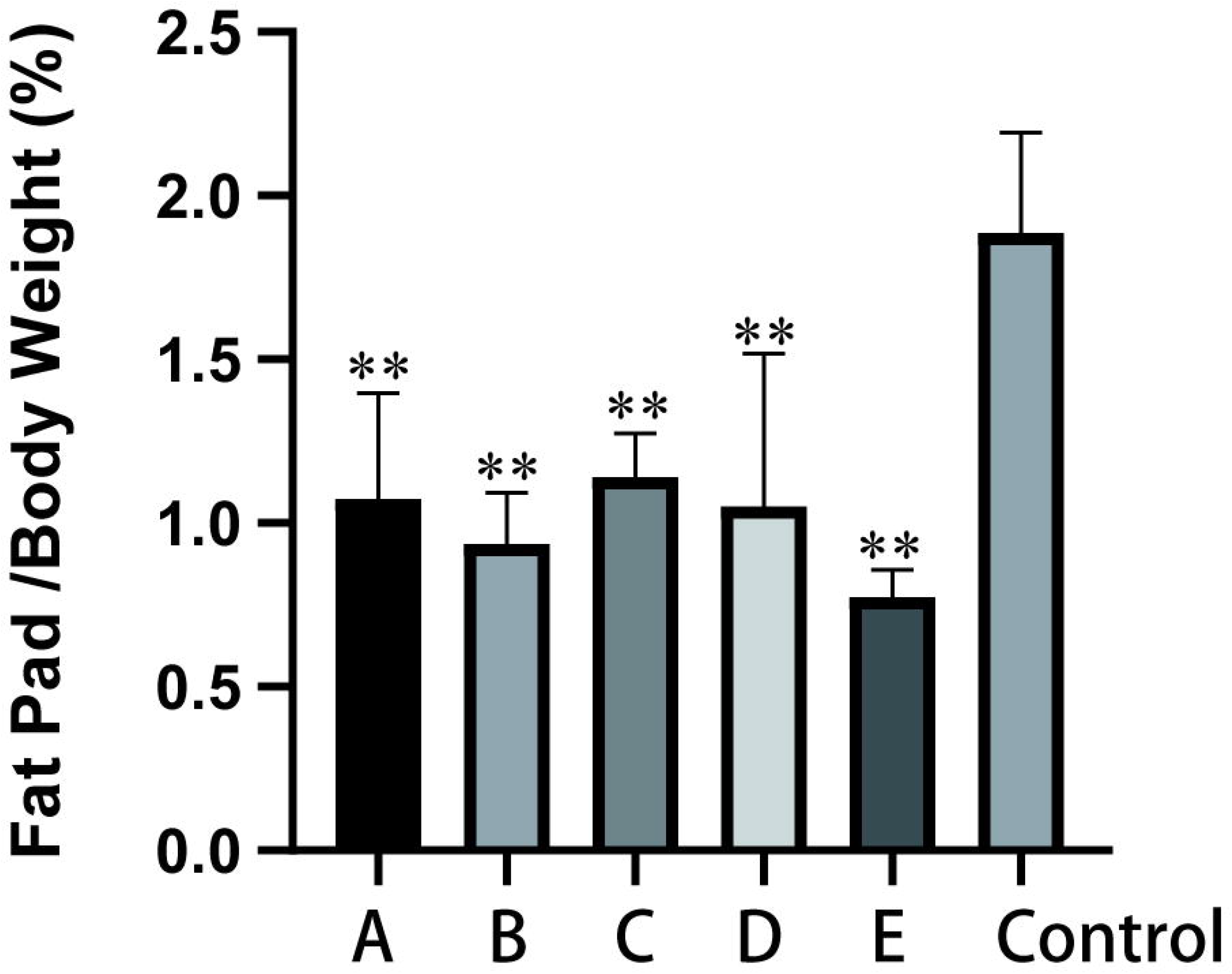

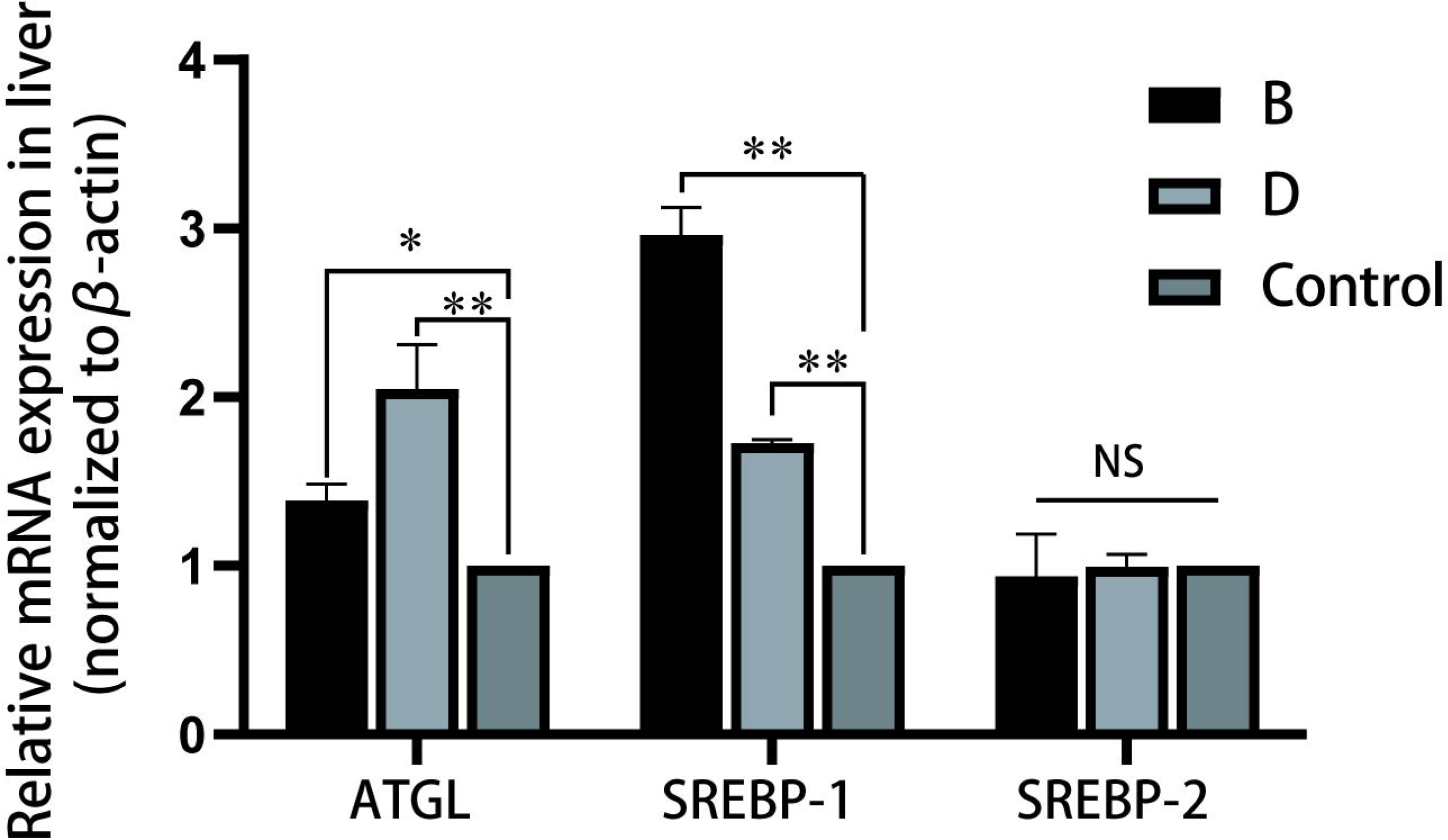

### 3.2. Jejunum morphology

In the poultry industry, the wide use of antibiotics usually results in a thinner intestine wall, smaller total villus area, shorter villus height, and crypt depth (Miles et al. 2006). In this study, all birds were purchased from a local farm (90 days), with no different treatment from the local commercial birds. The jejunum morphology analysis was performed in week 8 (**Table 4; Fig. 3**). The change found in the jejunum resulting from dietary supplements was significant as compared to control (*P*<0.01). The intestine wall thickness was significantly increased in the IPFF groups: approximately 79 μm, thicker than other groups (*P*<0.01). The villus height of group A, D, and E was approximately 800 μm, and achieved 908 μm in group B, suggesting that the intestinal villi growth were activated (Afsharmanesh and Sadaghi 2013). Group A and B showed a clear enhancement on the intestine wall, villus height, and crypt depth. Meanwhile, although the villus height and crypt depth showed no significant difference compared to the control group, the intestine wall was thicker in group C, suggesting that the IPFF additive was beneficial for the health of the intestine wall. In addition, a higher mucosal growth which required more energy for its performance (Awad et al. 2011), and so we assumed that the higher villus height and crypt depth in group B might be associate with the increased rate of tissue turnover to renew villi (Mehri et al. 2015).

**Table 4.** Jejunum Morphology Analysis of Laying Quails Fed a Basal Diet, IPFF Added Diet or IPR Added Diet.

|  | IPFF |  |  | IPR |  | Control | SEM | P |
| --- | --- | --- | --- | --- | --- | --- | --- | --- |
|  | A<br>(5%) | B<br>(1%) | C<br>(0.5%) | D<br>(5%) | E<br>(1%) |  |  |  |
| Intestine wall thickness(μm) | 78.5<sup>a</sup> | 79.2<sup>a</sup> | 76.9<sup>a</sup> | 43.7<sup>b</sup> | 43.4<sup>b</sup> | 41.3<sup>b</sup> | 6.09 | <0.01 |
| villous height (μm) | 835<sup>b</sup> | 908<sup>a</sup> | 575<sup>d</sup> | 818<sup>bc</sup> | 801<sup>c</sup> | 610<sup>d</sup> | 26.9 | <0.01 |
| Crypt depth (μm) | 112.8<sup>b</sup> | 189.5<sup>a</sup> | 94.7<sup>c</sup> | 72.9<sup>d</sup> | 77.0<sup>d</sup> | 91.5<sup>c</sup> | 4.35 | <0.01 |
| V/C<sup>a</sup> | 7.40<sup>b</sup> | 4.79<sup>d</sup> | 6.07<sup>c</sup> | 11.22<sup>a</sup> | 10.41<sup>a</sup> | 6.67<sup>b</sup> | 0.50 | <0.01 |

N=9. <sup>a</sup>. V/C means the ratio of villous length/crypt depth. **A**: Basal diet supplemented with 5% IPFF, **B**: basal diet supplemented with 1% IPFF, **C**: basal diet supplemented with 0.5% IPFF, **D**: basal diet supplemented with 5% IPR, **E**: basal diet supplemented with 1% IPR, **Control**: Control group (base diet); Superscripts **a, b, c, d, e** Means within the same superscripts without the same row are significantly different (P<0.05).

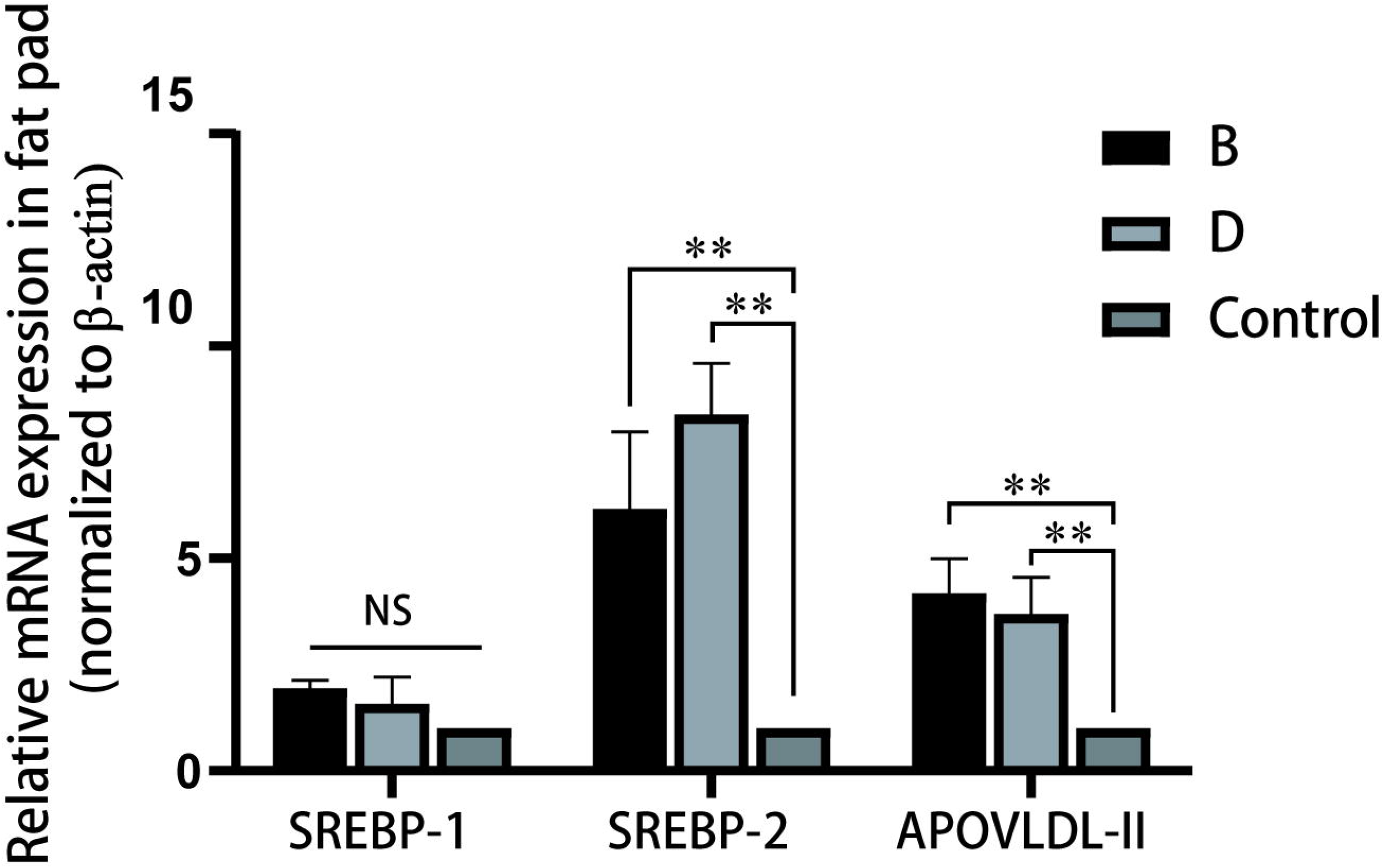

### 3.3. Yolk cholesterol and fatty acids

Cholesterol is essential for many physiological functions, including as a raw material for yolk formation and egg production (Schroeder et al. 1991; Simons and Ikonen 2000; Naviglio et al. 2012). However, high cholesterol intake must be avoided in our diet, because the excess of cholesterol is related to a range of diseases such as inflammation, diabetes, and cancer (Oliveira et al. 2011; Tajima et al. 2014; Wang X et al. 2019). Lower cholesterol egg was helpful to control cholesterol intake, thus is commonly considered as the superior quality (Zhong et al. 2019).

The determination of yolk cholesterol and yolk triglycerides is shown in **Table 5**. In this study, as compared to control, all groups showed no effects on triglycerides. The decrease in yolk cholesterol was observed in group B, C and E as compared to control (*P*<0.01), while the 5% groups (A and D) have no significant change. And the most significant yolk cholesterol decline was observed in the group B. These results indicate that the 1% addition of IPFF or IPR was the most effective on the yolk cholesterol regulation.

**Table 5.**
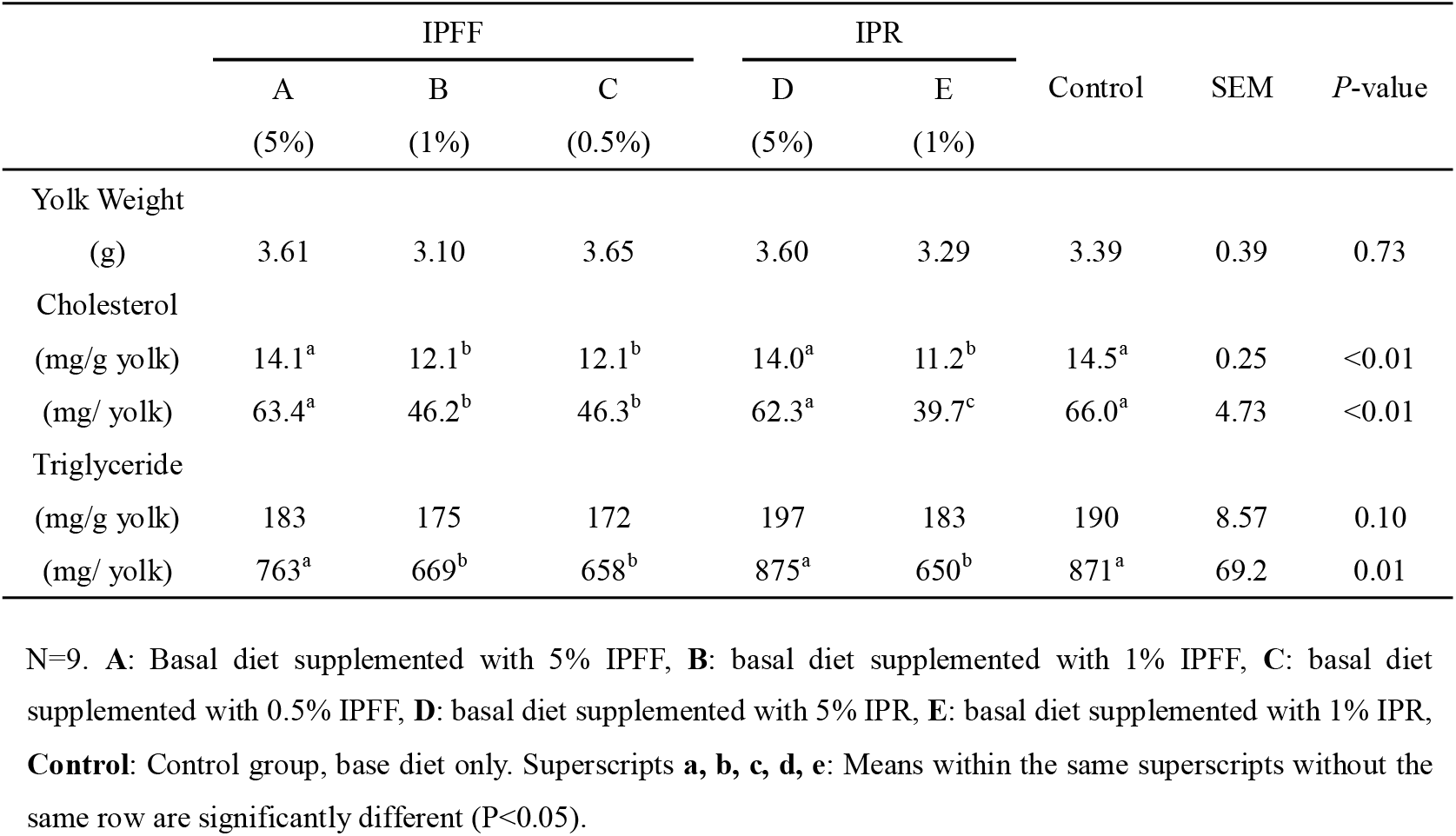
Yolk Cholesterol and Triglyceride Content of Laying Quails Fed a Basal Diet, IPFF Added Diet or IPR Added Diet.

|  | IPFF |  |  | IPR |  | Control | SEM | P-value |
| --- | --- | --- | --- | --- | --- | --- | --- | --- |
|  | A<br>(5%) | B<br>(1%) | C<br>(0.5%) | D<br>(5%) | E<br>(1%) |  |  |  |
| Yolk Weight |  |  |  |  |  |  |  |  |
| (g) | 3.61 | 3.10 | 3.65 | 3.60 | 3.29 | 3.39 | 0.39 | 0.73 |
| Cholesterol |  |  |  |  |  |  |  |  |
| (mg/g yolk) | 14.1<sup>a</sup> | 12.1<sup>b</sup> | 12.1<sup>b</sup> | 14.0<sup>a</sup> | 11.2<sup>b</sup> | 14.5<sup>a</sup> | 0.25 | <0.01 |
| (mg/ yolk) | 63.4<sup>a</sup> | 46.2<sup>b</sup> | 46.3<sup>b</sup> | 62.3<sup>a</sup> | 39.7<sup>c</sup> | 66.0<sup>a</sup> | 4.73 | <0.01 |
| Triglyceride |  |  |  |  |  |  |  |  |
| (mg/g yolk) | 183 | 175 | 172 | 197 | 183 | 190 | 8.57 | 0.10 |
| (mg/ yolk) | 763<sup>a</sup> | 669<sup>b</sup> | 658<sup>b</sup> | 875<sup>a</sup> | 650<sup>b</sup> | 871<sup>a</sup> | 69.2 | 0.01 |

N=9. **A:** Basal diet supplemented with 5% IPFF, **B:** basal diet supplemented with 1% IPFF, **C:** basal diet supplemented with 0.5% IPFF, **D:** basal diet supplemented with 5% IPR, **E:** basal diet supplemented with 1% IPR, **Control:** Control group, base diet only. Superscripts **a, b, c, d, e:** Means within the same superscripts without the same row are significantly different (P<0.05).

Ouyang et al found that the oil supplement given to hens was able to change the cholesterol and polyunsaturated fatty acid content in the yolk (Ouyang et al. 2004). Furthermore, in the research of Aydin et al, the dietary conjugated linoleic acid added to the hen’s diet significantly changed the fatty acid composition of the yolk (Aydin and Cook 2004). Indeed, both IPFF and IPR contained few lipids (**Table 1**): the conjugated linoleic acid in IPR oil was 68.64% and in IPFF oil was 72.21% (**Table 6**) which was in agreement with the results of Yang et al (Yang F-X et al. 2009). The composition of yolk fatty acids is shown in **Table 7**. However, the changes in fatty acids, including saturated acids and monounsaturated acids, were inconsistent with the change described in the study of Shinn et al (Shinn et al. 2015). Moreover, while the lipids content is different, the yolk cholesterol has no significant difference between groups B, C, and E. These results suggested that *Idesia polycarpa* oil might exert an effect upon the egg and birds, but it was not the main factor responsible for the changes in this study.

**Table 6.**
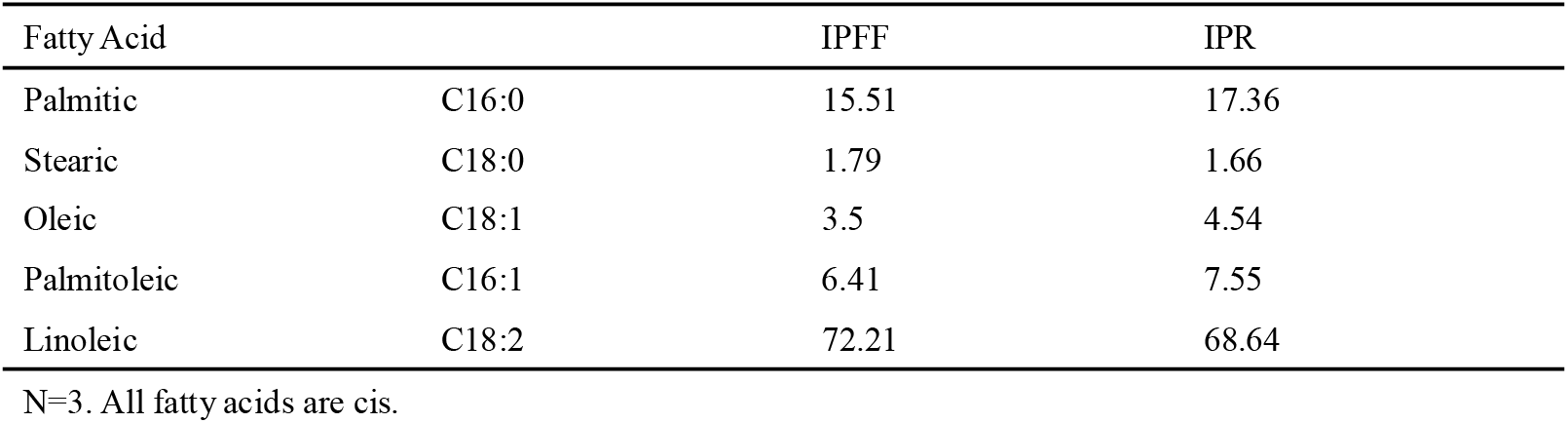
Fatty Acid Composition of IPFF Oil and IPR Oil (%)

**Table 7.** Fatty Acid Composition of Yolk Oil extracted from each groups.

| Fatty acid |  | Control | A | B | C | D | E |
| --- | --- | --- | --- | --- | --- | --- | --- |
| Myristic | C14:0 | 0.45<sup>b</sup> | 0.57<sup>a</sup> | 0.54<sup>a</sup> | 0.43<sup>b</sup> | 0.43<sup>b</sup> | 0.39<sup>b</sup> |
| Palmitic | C16:0 | 29.53 | 30.55 | 31.88 | 29.94 | 30.92 | 32.72 |
| Palmitoleic | C16:1;n-7 | 2.16<sup>c</sup> | 3.64<sup>a</sup> | 2.39<sup>bc</sup> | 1.75<sup>d</sup> | 2.82<sup>b</sup> | 2.20<sup>bc</sup> |
| Stearic | C18:0 | 12.8<sup>a</sup> | 11.6<sup>b</sup> | 10.5<sup>c</sup> | 10.1<sup>c</sup> | 9.23<sup>d</sup> | 10.6<sup>c</sup> |
| Oleic | C18:1;n-9 | 33.8 | 35.3 | 34.6 | 38.8 | 37.2 | 32.2 |
| Linoleic | C18:2;n-6 | 17.1<sup>b</sup> | 14.1<sup>c</sup> | 16.7<sup>b</sup> | 15.8<sup>b</sup> | 16.8<sup>b</sup> | 18.2<sup>a</sup> |
| α-Linolenic | C18:3;n-3 | 0.22<sup>a</sup> | 0.21<sup>a</sup> | 0.25<sup>a</sup> | 0.17<sup>b</sup> | 0.17<sup>b</sup> | 0.22<sup>a</sup> |
| Arachidonic | C20:4;n-6 | 2.56<sup>a</sup> | 2.38<sup>a</sup> | 1.88<sup>b</sup> | 1.84<sup>b</sup> | 1.65<sup>b</sup> | 1.64<sup>b</sup> |
| Eicosapentaenoic | C20:5;n-3 | 0.27<sup>b</sup> | 0.37<sup>a</sup> | 0.29<sup>ab</sup> | 0.23<sup>b</sup> | 0.14<sup>c</sup> | 0.25<sup>b</sup> |
| Docosahexaenoic | C22:6;n-3 | 0.89<sup>b</sup> | 1.02<sup>a</sup> | 0.76<sup>c</sup> | 0.68<sup>c</sup> | 0.51<sup>d</sup> | 0.48<sup>d</sup> |
| SFA |  | 42.8<sup>a</sup> | 42.7<sup>a</sup> | 42.9<sup>a</sup> | 40.5<sup>b</sup> | 40.6<sup>b</sup> | 43.7<sup>a</sup> |
| MUFA |  | 36.0<sup>c</sup> | 38.9<sup>b</sup> | 37.0<sup>bc</sup> | 40.6<sup>a</sup> | 40.0<sup>a</sup> | 34.4<sup>d</sup> |
| PUFA |  | 21.0<sup>a</sup> | 18.1<sup>d</sup> | 19.8<sup>b</sup> | 18.7<sup>c</sup> | 19.3<sup>b</sup> | 20.8<sup>ab</sup> |
| n-3PUFA |  | 1.38<sup>b</sup> | 1.60<sup>a</sup> | 1.30<sup>b</sup> | 1.08<sup>c</sup> | 0.82<sup>d</sup> | 0.95<sup>cd</sup> |
| n-6PUFA |  | 19.6<sup>a</sup> | 16.5<sup>d</sup> | 18.5<sup>b</sup> | 17.6<sup>c</sup> | 18.4<sup>b</sup> | 19.9<sup>a</sup> |
| n6/n3 |  | 14.2<sup>d</sup> | 10.3<sup>e</sup> | 14.3<sup>d</sup> | 16.3<sup>c</sup> | 22.5<sup>a</sup> | 20.9<sup>b</sup> |

N=3. **A:** Basal diet supplemented with 5% IPFF, **B:** basal diet supplemented with 1% IPFF, **C:** basal diet supplemented with 0.5% IPFF, **D:** basal diet supplemented with 5% IPR, **E:** basal diet supplemented with 1% IPR, **Control:** Control group (base diet); Superscripts **a, b, c, d, e** Means within the same superscripts without the same row are significantly different (P<0.05). All fatty acids are cis.

A lower n6/n3 ratio is good as the guidelines of WHO regarding healthy food for humans (Kris-Etherton et al. 2007). As shown in **Table 7**, the n6/n3 ratio was significantly higher than the control in groups D and E (*P*<0.01). Conversely, as compared to control, the decrease of n6/n3 ratio in group A was significant (*P*<0.01), with the increased n3 (EPA and DHA) and decreased n6 content that was more beneficial for the absorption of unsaturated fatty acid (Kris-Etherton et al. 2007).

### 3.4. Egg production and quality

The effect of IPFF or IPR dietary supplements on quail eggs was measured (**Table 8**). The results showed that the different dietary supplements produced no effect on egg or shell morphology. The yolk weight, shell thickness, Haught unit, and diameter ratio were similar among the groups. The yolk weight was approximately 0.29-0.33% of the total weight of the egg. Shell thickness was 0.16-0.18 mm, Haught unit was 79-84, and the diameter ratio was 0.77-0.82.

**Table 8.** Egg Quality of Laying Quails Fed a Basal Diet, IPFF Added Diet or IPR Added Diet.

|  | IPFF |  |  | IPR |  | Control | SEM | P-VALUE |
| --- | --- | --- | --- | --- | --- | --- | --- | --- |
|  | A<br>(5%) | B<br>(1%) | C<br>(0.5%) | D<br>(5%) | E<br>(1%) |  |  |  |
| Yolk/Egg<br>(in weight %) |  |  |  |  |  |  |  |  |
| initial | 0.30 | 0.30 | 0.31 | 0.31 | 0.30 | 0.31 | 0.187 | 0.980 |
| last week | 0.31 | 0.30 | 0.32 | 0.33 | 0.31 | 0.29 | 0.029 | 0.935 |
| average | 0.31 | 0.30 | 0.31 | 0.32 | 0.31 | 0.30 | 0.119 | 0.754 |
| Shell Thickness(mm) |  |  |  |  |  |  |  |  |
| Initial | 0.16 | 0.16 | 0.16 | 0.16 | 0.17 | 0.16 | 0.007 | 0.785 |
| Last week | 0.17 | 0.18 | 0.18 | 0.19 | 0.18 | 0.18 | 0.008 | 0.285 |
| Average | 0.18 | 0.18 | 0.18 | 0.18 | 0.18 | 0.17 | 0.008 | 0.563 |
| Haugh Unit |  |  |  |  |  |  |  |  |
| Initial | 80.59 | 79.84 | 78.12 | 80.75 | 79.59 | 79.15 | 2.760 | 0.827 |
| Last week | 79.25 | 81.18 | 80.42 | 79.86 | 80.42 | 80.78 | 1.881 | 0.924 |
| Average | 82.92 | 81.51 | 79.78 | 84.75 | 81.26 | 80.81 | 1.437 | 0.458 |
| Diameter Ratio |  |  |  |  |  |  |  |  |
| Initial | 0.77 | 0.80 | 0.78 | 0.82 | 0.79 | 0.80 | 0.346 | 0.805 |
| Last week | 0.77 | 0.78 | 0.78 | 0.78 | 0.78 | 0.79 | 0.082 | 0.285 |
| Average | 0.78 | 0.77 | 0.78 | 0.79 | 0.78 | 0.77 | 0.007 | 0.269 |

N=9. **Control**, Control group (base diet); **A**, basal diet supplemented with 5% IPFF; **B**, basal diet supplemented with 1% IPFF; **C**, basal diet supplemented with 0.5% IPFF; **D**, basal diet supplemented with 5% IPR; **E**, basal diet supplemented with 1% IPR.
**Initial**: Mean of week 0 to 2; **Last week**: Mean of week 8; **Average**: Mean of week 2 to 8.

Low dose IPFF dietary supplementation enhanced the egg production capacity (**Fig. 4**). Group C kept a steady increase as compared to the control. The egg production of the IPFF groups was increased during the whole experiment period, compared to the control (**Table 9**). The egg production increased in the IPFF groups at different levels but decreased in the IPR added groups. Group B (1% IPFF added) achieved the highest production in the last week of phase 2, since it was 90.2% on average, approximately 15% above the control. Furthermore, the results of phase 3 show that the layer quails production was still affected by the previous IPFF or IPR supplementation. Likewise, the IPFF and IPR supplementation were changing the egg mass effectively. The average egg mass in group B was 9.77, which was significantly higher than in other groups. On the contrary, the egg mass decreased in IPR groups: group D (5% IPR added) exhibited a significant dose-related production decrease which particularly in the last week as compared to control (week 8; *P*<0.01).

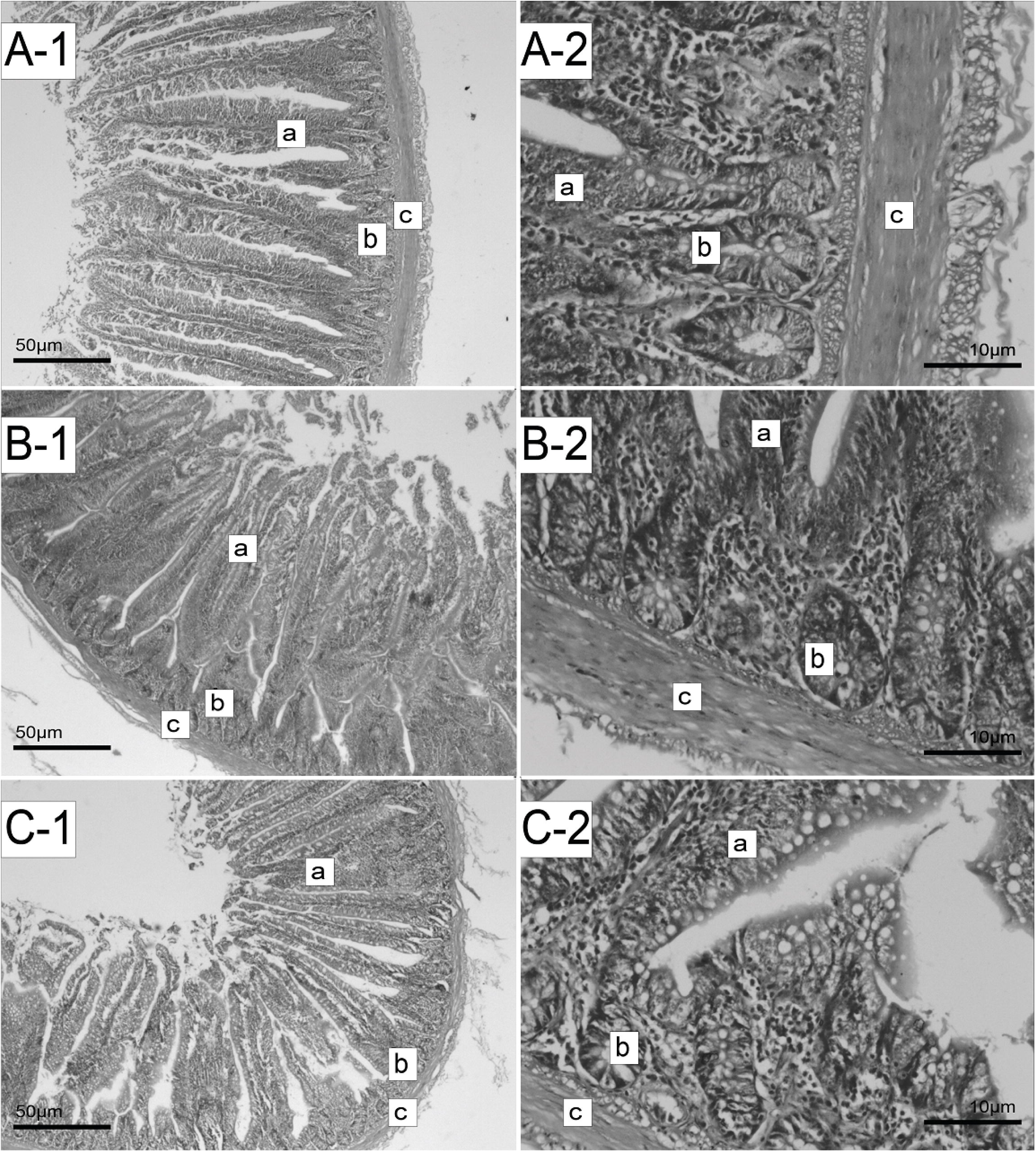

**Table 9.** Egg Mass and Egg Production of Laying Quails Fed a Basal Diet, IPFF Added Diet or IPR Added Diet.

|  | IPFF |  |  | IPR |  | Control | SEM | *P*-value |
| --- | --- | --- | --- | --- | --- | --- | --- | --- |
|  | A | B | C | D | E |  |  |  |
|  | (5%) | (1%) | (0.5%) | (5%) | (1%) |  |  |  |
| Egg mass (g/d/bird) |  |  |  |  |  |  |  |  |
| Initial | 9.44<sup>a</sup> | 9.41<sup>a</sup> | 9.25<sup>a</sup> | 9.31<sup>a</sup> | 9.14<sup>a</sup> | 9.18<sup>a</sup> | 0.30 | 0.88 |
| Last week | 9.18<sup>b</sup> | 9.31<sup>ab</sup> | 9.58<sup>a</sup> | 6.04<sup>d</sup> | 7.82<sup>c</sup> | 9.27<sup>ab</sup> | 0.30 | <0.01 |
| Average | 9.41<sup>a</sup> | 9.77<sup>a</sup> | 9.40<sup>a</sup> | 6.06<sup>d</sup> | 7.86<sup>c</sup> | 8.87<sup>b</sup> | 0.38 | <0.01 |
| Phase 3 | 8.19<sup>b</sup> | 9.01<sup>a</sup> | 8.63<sup>ab</sup> | 6.24<sup>d</sup> | 7.13<sup>c</sup> | 8.37<sup>ab</sup> | 0.32 | <0.01 |
| Egg production (%) |  |  |  |  |  |  |  |  |
| Initial | 81.0<sup>a</sup> | 81.3<sup>a</sup> | 81.3<sup>a</sup> | 80.7<sup>a</sup> | 80.7<sup>a</sup> | 79.3<sup>a</sup> | 2.57 | 0.97 |
| Last week | 80.1<sup>b</sup> | 90.2<sup>a</sup> | 83.8<sup>b</sup> | 56.3<sup>e</sup> | 68.1<sup>d</sup> | 78.4<sup>c</sup> | 2.91 | <0.01 |
| Average | 82.2<sup>a</sup> | 87.7<sup>a</sup> | 81.7<sup>a</sup> | 62.4<sup>c</sup> | 72.3<sup>b</sup> | 76.0<sup>b</sup> | 3.15 | <0.01 |
| Phase 3 | 70.3<sup>b</sup> | 81.5<sup>a</sup> | 75.8<sup>ab</sup> | 59.2<sup>c</sup> | 63.5<sup>c</sup> | 72.3<sup>b</sup> | 2.92 | <0.01 |

N=9. **A:** Basal diet supplemented with 5% IPFF, **B:** basal diet supplemented with 1% IPFF, **C:** basal diet supplemented with 0.5% IPFF, **D:** basal diet supplemented with 5% IPR, **E:** basal diet supplemented with 1% IPR, **Control:** control group (base diet); Superscripts **a, b, c, d, e** Means within the same superscripts without the same row are significantly different (P<0.05). **Initial:** Mean of week 0 to 2; **Last week:** Mean of week 8; **Average:** Mean of week 8 to 10; **Phase 3:** Mean of week 10 to 11.

Compared to the 5% IPFF group, lower IPFF amount (1% and 0.5%) added groups were more effective to promote egg production and egg mass (*P*<0.01). Furthermore, the result also suggested that most of the potential anti-nutritional factors of IPR that are harmful to nutrients absorption could be alleviated by solid fermentation.

The additive amount of 1% is the most efficient in egg quantity, while we noticed that the egg mass of group B was decreased as the experiment went on. The egg mass of group B was significantly higher than in other groups but lower than that in group C and similar to control in the last week (week 8). It is well known that lipid mobilization from the adipose tissue is one of the primary sources that support the onset of egg production. As a result of lipid mobilization, the increased fat pad is the most intuitive reflection of raw material supply for egg production (Yang S et al. 2013). In this study, the interclavicular fat pat weight was significant decreased in both IPFF added groups and IPR added groups. However, the ATGL and SREBP-1 were up-regulated in the fat pad which means the enhanced synthesis of the lipids (**Fig. 2**). Meanwhile, up-regulated SREBP-2 and APOVLDL-II in the liver suggested that the synthesis and transportation of cholesterol were enhanced. The enhanced lipids metabolism could supply abundant raw materials for body physiological processes. And egg formation is the most important process at this stage. Besides, described jejunum morphology results indicated that the birds of group B had a better intestine health situation which made better digestion and absorption (**Fig. 5**).

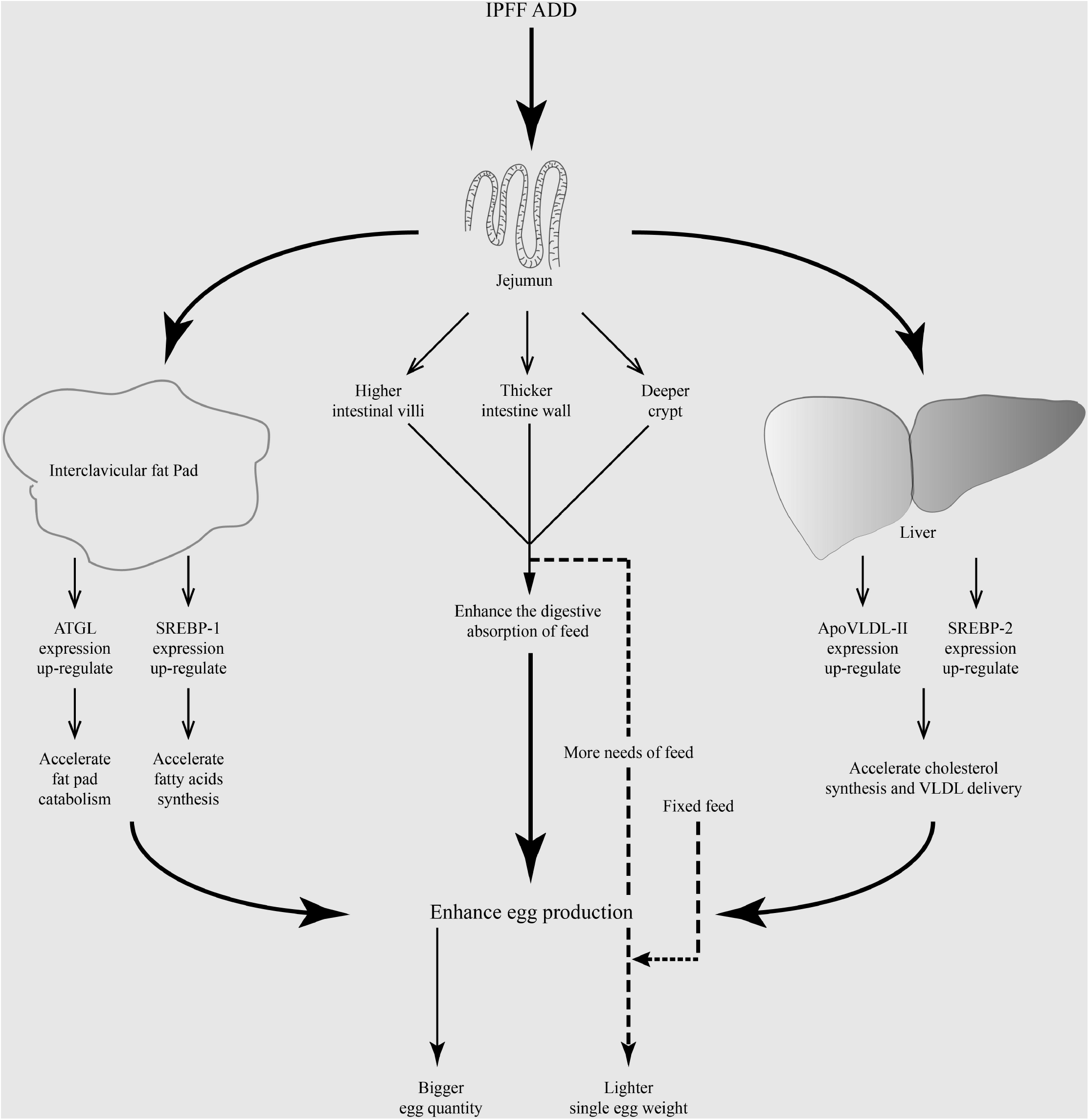

However, the daily feeding amount was strictly controlled to 20 g per bird to observe the change of the egg production more indicative. IPFF has promoted the quantity of eggs, but the eggs were lighter due to the fixed diet, just as the results showed in group B. These results indicated that while the raw-material supplement which supplies for the egg production was enhanced in IPFF groups, but with the improvement of product performance, the fixed diet cannot meet the requirement of gradually strengthened production performance. The egg mass of group B was picked up in phase 3, this is also bearing this opinion out. Moreover, the same tendency of gene expression was found in both groups B and D, combined with the change of fat pad. And as such, we hold the opinion that the bio-active substances which existed in *Idesia polycarpa* residues were able to affect the egg formation and production of laying quails. However, the key substance which plays an essential role still needs to be confirmed in subsequent studies. And the taste, by the way, after a small scope test (20 persons) we found, the eggs obtained from IPFF added groups were still as delicious as a commercial egg.

## 4. Conclusion

Overall, our research showed that IPFF could be a good feeding additive for laying quails which help to increase egg production and quality. 1% added was the most efficient dosage of IPFF that not only for the enhance of egg production but also for the decrease of yolk cholesterol. Meanwhile, although 5% IPFF added had no significant effect on egg production, but it helps to a higher EPA and DHA content and a better n3/n6 ratio. Furthermore, IPFF was helpful for the health of the quail’s intestine and reduced the potential negative effect of industrial feeding. To our knowledge, this study is the first time showing the application of *Idesia polycarpa* residues in poultry feeding. Moreover, we found a new method to transform the “useless” *Idesia polycarpa* residues into valuable agriculture products.

## Abbreviations

*Idesia polycarpa, Idesia polycarpa Maxim*. var. *vestita Diels*;

IPR: *Idesia polycarpa* residue
IPFF: *Idesia polycarpa* residue fermented feed
PUFA: Polyunsaturated fatty acid

